# Targeted Vibration-Induced Necrosis in Cancer Cells using Paramagnetic Microrobots

**DOI:** 10.1101/2022.10.19.512945

**Authors:** Sudipta Mallick, Max Sokolich, David Rivas, Sambeeta Das

**Author notes:** Corresponding Author: Prof. Sambeeta Das.

## Abstract

Therapeutic delivery of anti-cancer drugs is a major goal of modern medicine. In particular, microrobots (MRs) have recently been studied for their ability to navigate difficult-to-reach regions in the human body to deliver therapeutics for microscopically localized interventions. However, the control of individual and swarms of MRs to precisely target localized cellular regions remains a significant challenge, preventing their applications as delivery systems in cancer research. In this study, magnetic MRs were used to target cancer cells and create localized magnetic oscillations which resulted in magnetolysis of cancer cells. The magnetic MRs were selectively steered towards Hepatocarcinoma cells (HepG2 cells) using our custom-built magnetic controller under a rotating magnetic field at different frequencies. After internalization of the microrobots by cancer cells, magnetic oscillation of varying dosages was applied to disrupt the internal structure of cancer cells which leads to subsequent cell death.

## Introduction

Material concept and design has undergone significant advancement in the biomedical field in order to achieve desired structures, properties, and functionalities ^1^. Recently, nature-inspired design-strategies have been adopted by various interdisciplinary research groups to develop smart and sustainable devices for medical and healthcare applications^2,3,4^. Microrobots (micro/nano-sized robots) are one such invention that are inspired by micro-organism such as bacteria, motor proteins and spermatozoa etc. to perform various biomedical tasks ^5^. Microrobots (MRs) are highly desirable because of their relatively large size which allows them to be more readily visualized under a microscope compared to nano-size robots ^6^. These MRs can be driven by various chemical (H_2_O_2_, H_2_O and NaBH_4_) and external energy fields (magnetic field, UV, ultrasound) through narrow and confined spaces with precise control^7^. Particularly, magnetic MRs are of great interest due to their ability to be actuated under the influence of a magnetic field allowing for the use of a custom-designed workspace with various motion control methods ^8^. Magnetically actuated MRs can be steered to different inaccessible parts of the body without resorting to invasive procedures. Moreover, diverse structural and surface properties of magnetic MRs offer potential uses in the field of delivery systems, tracking, and single cell manipulation ^8, 9^.

In a living organism, every single activity is a function of all the biochemical reactions that happen at a cellular level. These biochemical reactions are often a collection of all the synchronized processes that are carried out at a subcellular level inside the cells ^10^. Any alteration in these highly specialized pathways can corrupt specific or multiple biological signals which often lead to disease onset ^11^. Cancer is the simplest example of a dysfunctional mechanism inside a cell that causes uncontrolled growth of the cells ^12^. Birth defects are another example where abnormalities in a single cell lead to impaired development of the fetus ^13^. Furthermore, correlations between abrupt changes in intracellular electric current across the cell and cardiac arrhythmia are also evident from previous research ^14^. Likewise, dysfunctional cellular mechanisms are the apparent cause of neurodegenerative diseases and diabetes as well ^15,16^. Therefore, targeting these subcellular pathways or microstructures have been important in drug discovery and development ^17^.

Recent reports have demonstrated that the application of a low frequency magnetic field (LFMF) effectively inhibits cancer cell proliferation and triggers apoptosis ^18^. Different magnetic particles like magnetic disks, iron microparticles, carbon nanotubes and magnetized-silica spindle-shaped particles were used to generate low-frequency magnetic field that kills cancer cells (magnetolysis) ^19,20,21,22^. While LFMF is known to inhibit cell proliferation and induce apoptosis by metabolic shift and affecting gene expression, respectively, ^23,24^ these reports have reported different mechanisms of cell death either by direct cell membrane damage or mechanical stress-induced apoptosis ^22,25^. Although these studies have demonstrated the magnetolysis effect of magnetic particles on cancer cells in different experimental set up, effect of LFMF and mechanism of cell death in a single cell level is yet to be determined.

In this present study, we use magnetically oscillating microrobots to induce cell lysis. The MRs were internalized by the cells, which does not result in toxic effects. Cancer cells with internalized MRs were then aligned in the direction of magnetic fields and oscillated via fast rotating magnetic fields in the xy plane. These results show that magnetic microrobots can be used as an effective means to kill cancer cells by a straightforward application of an alternating magnetic field.

## Results

### Control and transport of MRs

Experiments were carried out using our magnetic system described in Fig. 1. Paramagnetic (4,7μm) magnetic MRs were transported from one assigned place to another using our custom-built magnetic controller. Helmholtz coils were used to generate a rotating magnetic field that allows these MRs to roll along the surface of the substrate. Moreover, magnetic oscillation at various frequencies was created by switching (left or right) to different magnetic coils in the controller. Our unique and portable magnetic controller system was able to perform two different tasks: MRs were moved towards a specific location and subsequently vibrated.

**Figure-1:**
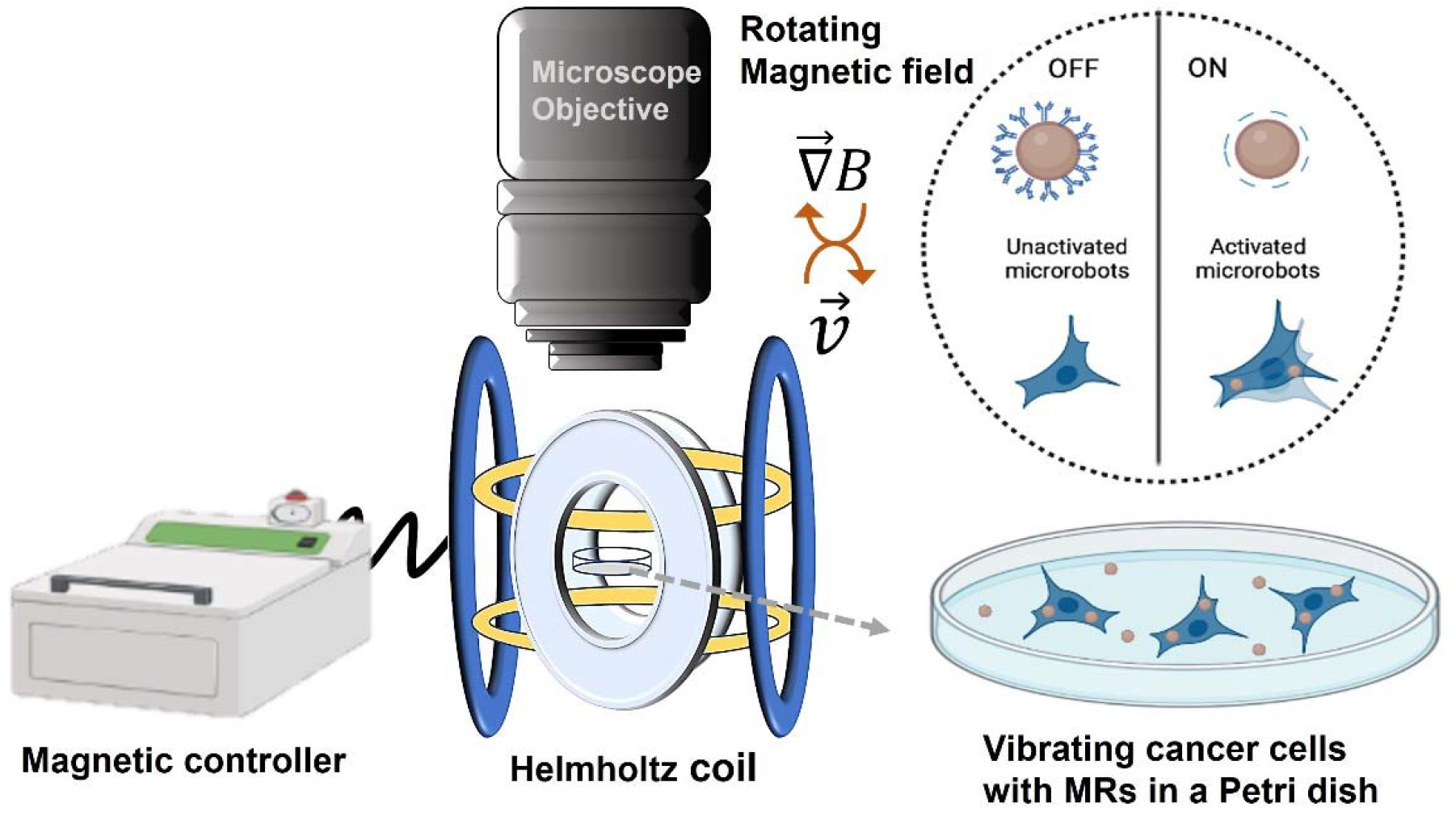
Schematic representation of killing cancer cells via magnetic oscillation.

### Cyto-compatibility of MRs

Biocompatibility is one of the most important factors in microrobot design. Apart from being non-toxic, MRs should not release any chemicals or by-products that affects results during manipulation. Here, paramagnetic beads are made of nontoxic polystyrene (figure 2a). Fluorescent MRs were used for all cell experiments to visualize these particles without further modification. Cells were incubated with the MRs for 24hrs, and changes in cell morphology was monitored using an optical and fluorescence microscope. The cell monolayer was intact without any toxic effects (Fig. 2b and c). Cells were then sorted to separate cell population with MRs and culture for 24hrs. As shown in Fig 3a and b, cells were adhered to the petridish without any dead suspended cells. Quantitative analysis was also consistent with the data showing negligible cell death of 3% when compared with untreated cells (Fig. 3 c).

**Figure-2:**
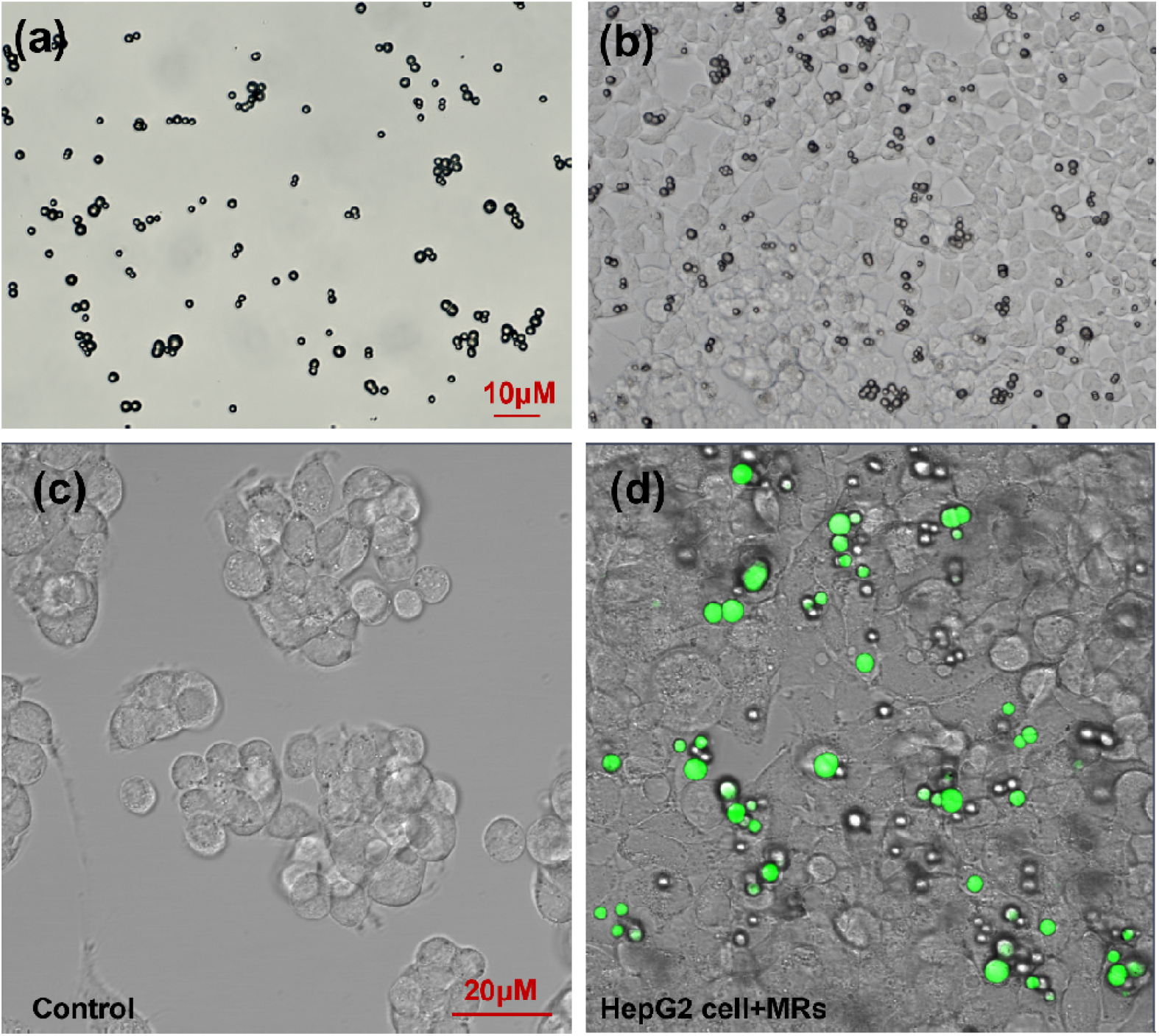
(a) Microscopic image of MRs, (b) adherent HepG2 cells with MRs and (c) Fluorescence image of the MRs (green) inside the cells.

**Figure-3:**
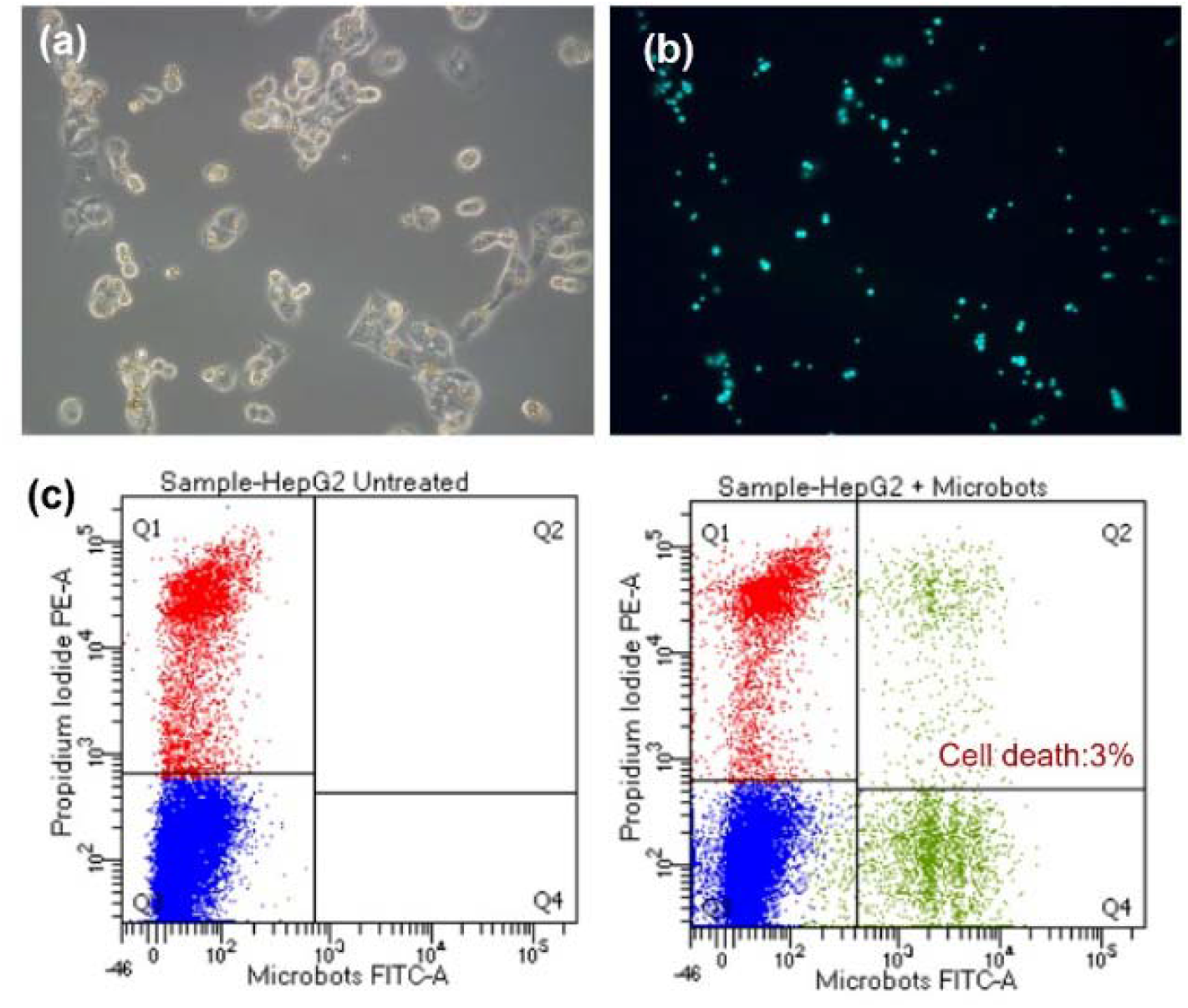
Cytocompatibility of MRs. HepG2 Cell treated with MRs (a) bright field image and (b) corresponding fluorescence image. Flow cytometry data showing cell death in (c) untreated and microrobot-treated cells.

### Cell-internalization of MRs

The MRs were completely internalized by the HepG2 cells after 24hrs. There were multiple MRs observed inside the cells showing high affinity of these polystyrene beads towards the cells (Fig. 2 d). Cell internalization was quantified using flow cytometry and 16.6% cells were with MRs when compared with control group (Fig 4). HepG2 cells with a broad range of intensities demonstrating inequal distribution of MRs insides the cells. This difference in cell internalization was presumably due to the cluster forming growth pattern of HepG2 cells that limits the surface area of interaction between cells and MRs.

**Figure-4:**
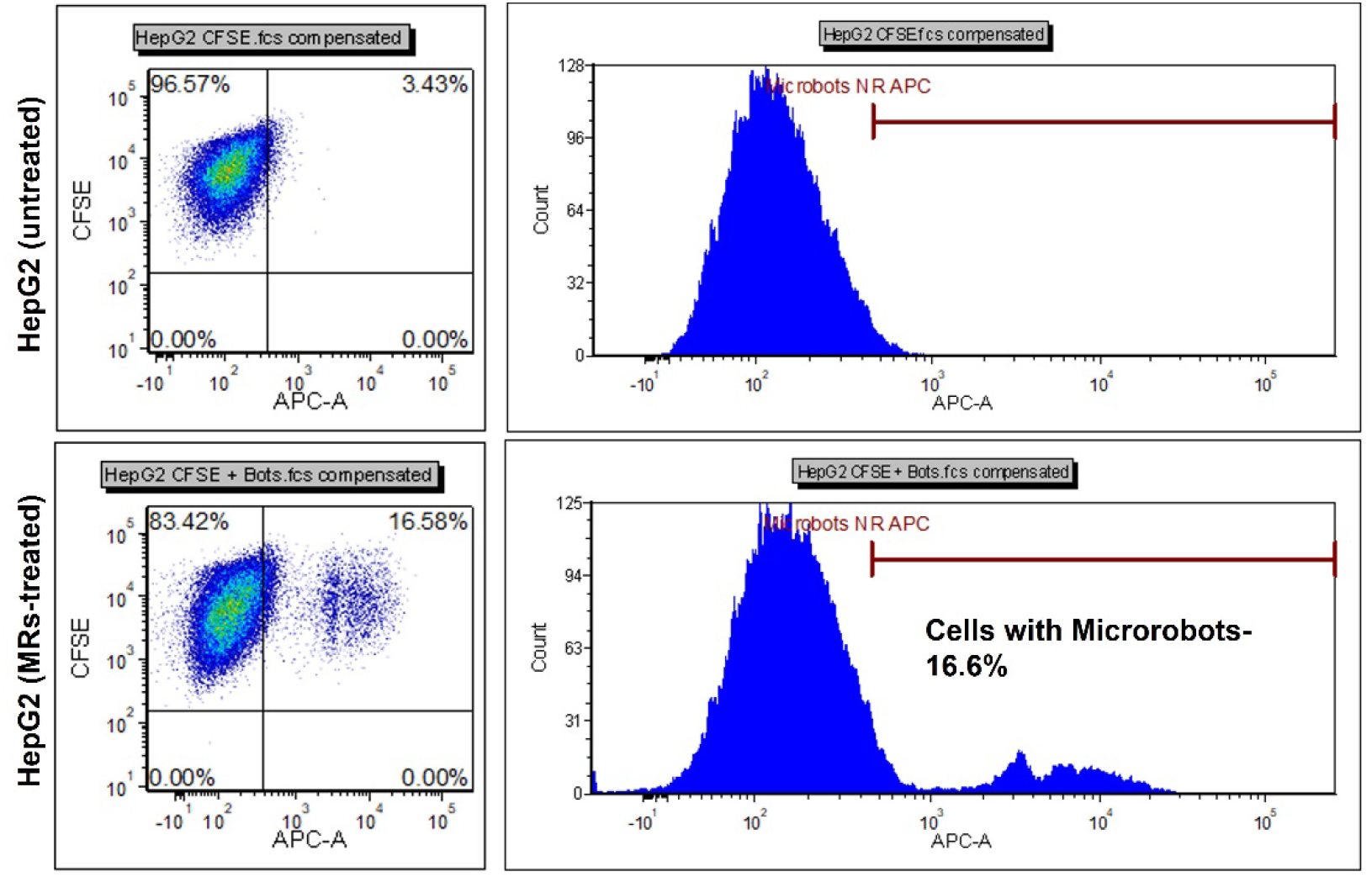
Cell internalization of MRs. Flow cytometry data showing cell internalization of MRs in HepG2 cells.

### Cell transport and targeted Vibration delivery

Targeting intracellular structures or organelles using MRs provides an opportunity to potentially alter or interfere with the cell’s biological functionality. We envisaged to apply intracellular vibration approach to disrupt the internal structure of cancer cells via magnetic oscillation of the MRs which ultimately leads to cell death. Once the MRs are internalized, we applied an oscillating square wave of predefined frequency to opposite facing electromagnets. As a result, the electromagnets rapidly changed from an on state to an off state asynchronously thereby resulting in a magnetic oscillation of the MRs. As shown in Fig. 5 and Vid. 1, the total angular displacement, Δ?, of the MRs between these cycles of field reversal depended on the applied frequency of the oscillating magnetic field. This is due to the rotational viscous drag on the microrobot which prevents it from fully aligning with the applied magnetic field prior to each field reversal. Results from this experiment showed that the HepG2 cells were responsive to oscillation and dead after 24hrs (Fig. 5). Vibration was applied for 30mins at two different frequencies (5 and 10 Hz) on HepG2 cells and increased cell death was observed just after one application (Fig. 6). Moreover, an interesting observation was found that cells with multiple number of MRs were dead due to the stronger oscillational force collectively generated by all MRs confirmed by fig. 5c where oscillatory strength is higher in cells with multiple MRs. This indicates a potential relationship between internal disruption and biochemical signaling. We assume the mechanism of internal disruption and cell death is caused by shear stress that is generated in the cytoplasm due to the magnetic oscillation. Chiew *et al*. have also reported similar findings that shows mechanical stress-induced cancer cell death using microparticles ^25, 26^. Moreover, propidium iodide staining during flow cytometry analysis also confirmed that membrane damage could be the possible cell destruction mechanism.

**Figure-5:**
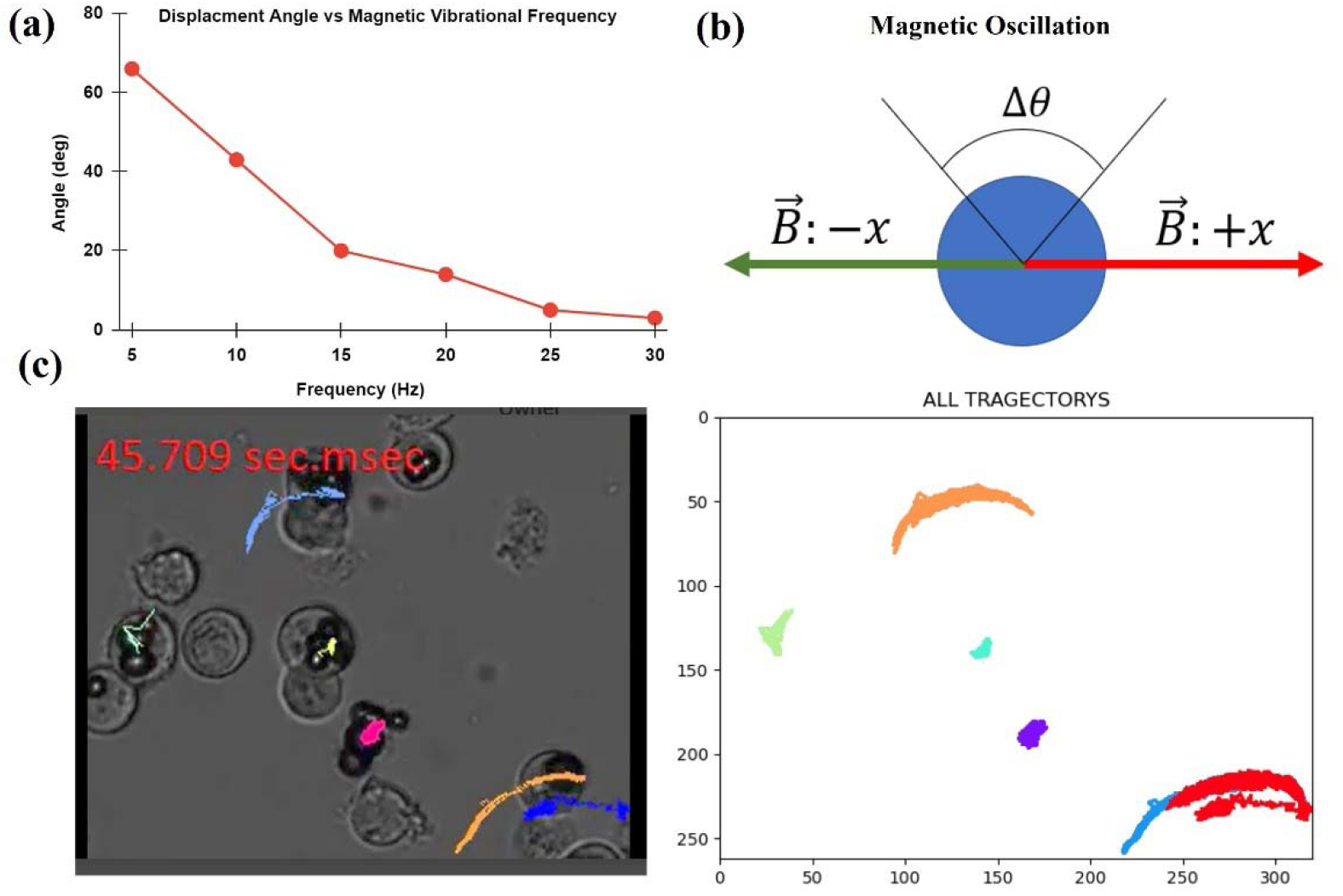
(a) Graph displaying displacement angle (Δθ) verses frequency and (b) a Schematic illustration of magnetic Oscillation. As the frequency at which each electromagnet is pulsed increases, the resulting angular displacement of the magnetic microrobot decreases. A frequency of 5 Hz results in the microrobot rotating approximately 60 degrees whereas a frequency of 30 Hz results in very small angular displacement. **(c)** Oscillating trajectory of Cells with MRs in different frequencies.

**Figure-6:**
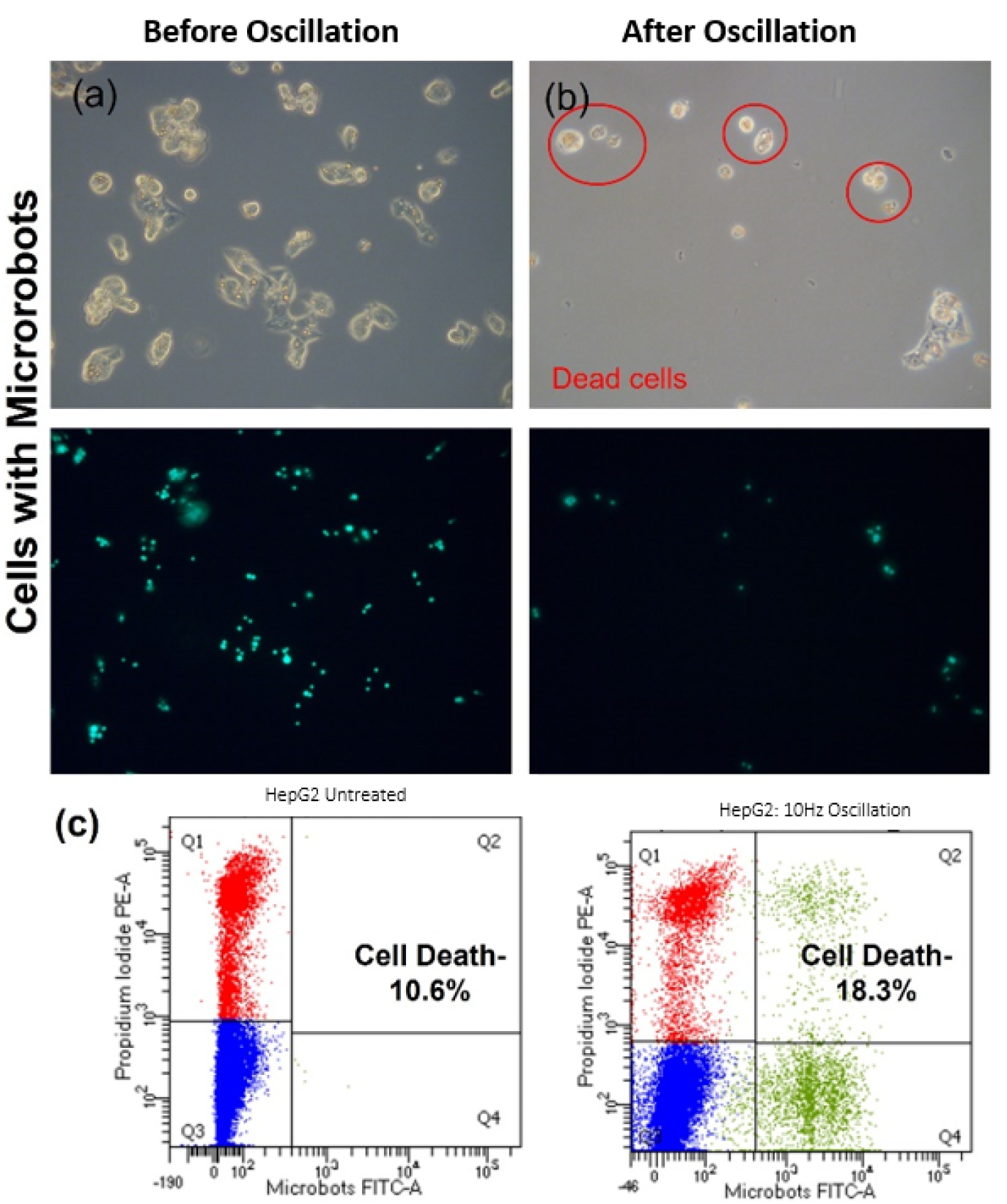
Cell death after oscillation. Fluorescence microscopic images of HepG2 cells (a) before and (b) after oscillation. Cells were trypsinzed and subjected to magnetic oscillation for 30mins and transferred to incubator. Cells with MRs were found dead after 24hrs (in red circles).

## Discussion and Future Directions

Our results demonstrate that low frequency mechanical vibration at 10 Hz promotes cell death in liver cancer cells. Previous research has shown that mechanical vibration of anisotropic magnetic nanoparticles attached to cell membranes can trigger apoptosis at subkilohertz frequencies ^27^. Our results demonstrate that low frequency mechanical vibration has the potential for selective cancer cell death. In addition, since an increase in temperature is unlikely to be elicited at such a low frequency, the cell death is most likely due to cell membrane or intracellular microstructure damaged-induced necrosis. Nevertheless, reports suggests that localized necrotic cell debris can enhance the effect of immunotherapy by evoking immune cells activation ^28^. Precise controllability over our magnetic system also offers us the ability to accurately navigate the robots to the complex locations in a cellular system.

The mechanisms underlying the effects of mechanical vibration on cell death are largely unknown. However, our results indicate that MRs can be an effective and non-invasive approach for analysis and investigation at the cellular level with improved accuracy and repeatability. Additionally, MRs can be effectively targeted to the desired location and serve as delivery systems for various biomedical applications. In the future, we plan to examine the effect of magnetic MRs induced vibrations on other cell and tissue types, whether cancerous or healthy ones, and in animal models. In addition, we would also study the underlying mechanism of vibration induced cell death. Our present study shows promising potential for magnetic oscillation-based therapeutic approaches for cancer and magnetic manipulation-based applications for better understanding of cell behavior and related biochemical outcomes, representing a new arena for cellular medicine.

## Materials and Methods

The MRs consisted of paramagnetic beads (diameter= 4.7 μm), which were purchased from Spherotech (Cat. No. FCM-4056-2).

## Magnetic experimental setup of MRs

Experiments were conducted on an Axioplan 2 upright microscope using an Axiovert 503 mono camera and an Axiovert 200 inverted microscope with a Amscope MU903-65 camera. Experiments on the Axioplan microscope utilized a custom-built magnetic control system which applied magnetic field strengths at the sample of approximately 3-7 mT. The magnetic system consists of four electromagnetic solenoids containing soft iron cores which are arranged in an array along the x and y axes, allowing for magnetic fields to be applied in any orientation in the horizontal plane. The strength and direction of the fields are controlled using custom Matlab code. The MATLAB program utilizes the Data Acquisition Toolbox and a Wireless gaming controller. The program connects to the controller through bluetooth and each button can be mapped to a certain digital output on a National Instruments DAQ USB 6351, and thus a specific electromagnet. Experiments performed on the axiovert microscope utilized hand-held permanent cylindrical magnets that produced magnetic field strengths at the sample of approximately 10-30 mT. Paramagnetic beads used in the experiments were fluorescent and had a diameter of 4.7 μm. The fluorescent microspheres were illuminated using an X-cite mini plus from excelitas technologies. Custom software was written in python to detect MRs and plot their trajectories.

### Cell culture and maintenance

Hepatocellular Carcinoma cells (HepG2) cells were gifted by Richard West (Associate Scientist at Flow Cytometry Core Facility). Cells were cultured in Dulbecco’s Modified Essential Medium (DMEM) and Ham’s F-12 (Gibco, BenchStable, USA) media with 5% CO_2_ and maintained at 37 °C in an incubator. All experiments were performed after third passage of cells.

### Assessment of Cytotoxicity and Cellular uptake of MRs

Cytotoxicity of MRs was evaluated in HepG2 cells. Cells were seeded (2×10^4^ cells/well) in a clear flat bottom 24-well plate (Costar, Corning, USA) and incubated in 1:1 mixture of Dulbecco’s Modified Essential Medium (DMEM) and Ham’s F-12 (Gibco, BenchStable, USA) media with 5% CO2 at 37 °C for 24hrs. Then, cells were treated with MRs (4.7μm size, Yellow, 1 mg/mL) and incubated for 24hrs. Cells were then imaged under optical microscope to check cell morphology. Flow cytometry was also performed to quantify exact percentage of cell death after propidium iodide (PI) staining. Cellular internalization of MRs was assessed by flow cytometry. Samples for cellular uptake were prepared as described by the aforementioned method in HepG2 cells. 1×10^5^ cells/well were seeded in a 6-well plate and incubated in DMEM/F-12 with 5% CO_2_ at 37 °C for 24hrs. Then, 20μL of 1% w/v MRs solution was added to the each well containing 2mL media. After 24hrs, cells were washed and stained with CFDA-SE for 15 mins and analyzed using a flow cytometer (BD FACS Aria llu).

### Cancer cell killing using magnetic oscillation

HepG2 cells were treated with MRs (100μg/mL) in a 6-well plate (Costar, Corning, USA) and incubated in DMEM/F-12 (Gibco, BenchStable, USA) media with 5% CO_2_ at 37 °C for 24hrs. After 24hrs, cells were washed and trypsinized to detach cells from the culture dish. Then, single HepG2 cells with internalized particles were subjected to magnetic oscillation with different frequency (frequency range: 5-10 Hz) in xy-plane for 30minutes.

## Acknowledgment

The authors gratefully acknowledge the late Richard West for his help with the cell lines. This project was supported by the Delaware INBRE program, with a grant from the National Institute of General Medical Sciences – NIGMS (P20 GM103446) from the National Institutes of Health and the State of Delaware. This work was also supported by NSF grant OIA2020973. This content is solely the responsibility of the authors and does not necessarily represent the official views of NIH.

## Data availability Statement

All relevant data are within the paper and its supporting information files.

