## Supplementary Video for "Targeted Vibration-Induced Necrosis in Cancer Cells using Paramagnetic Microrobots"

### Slide 1
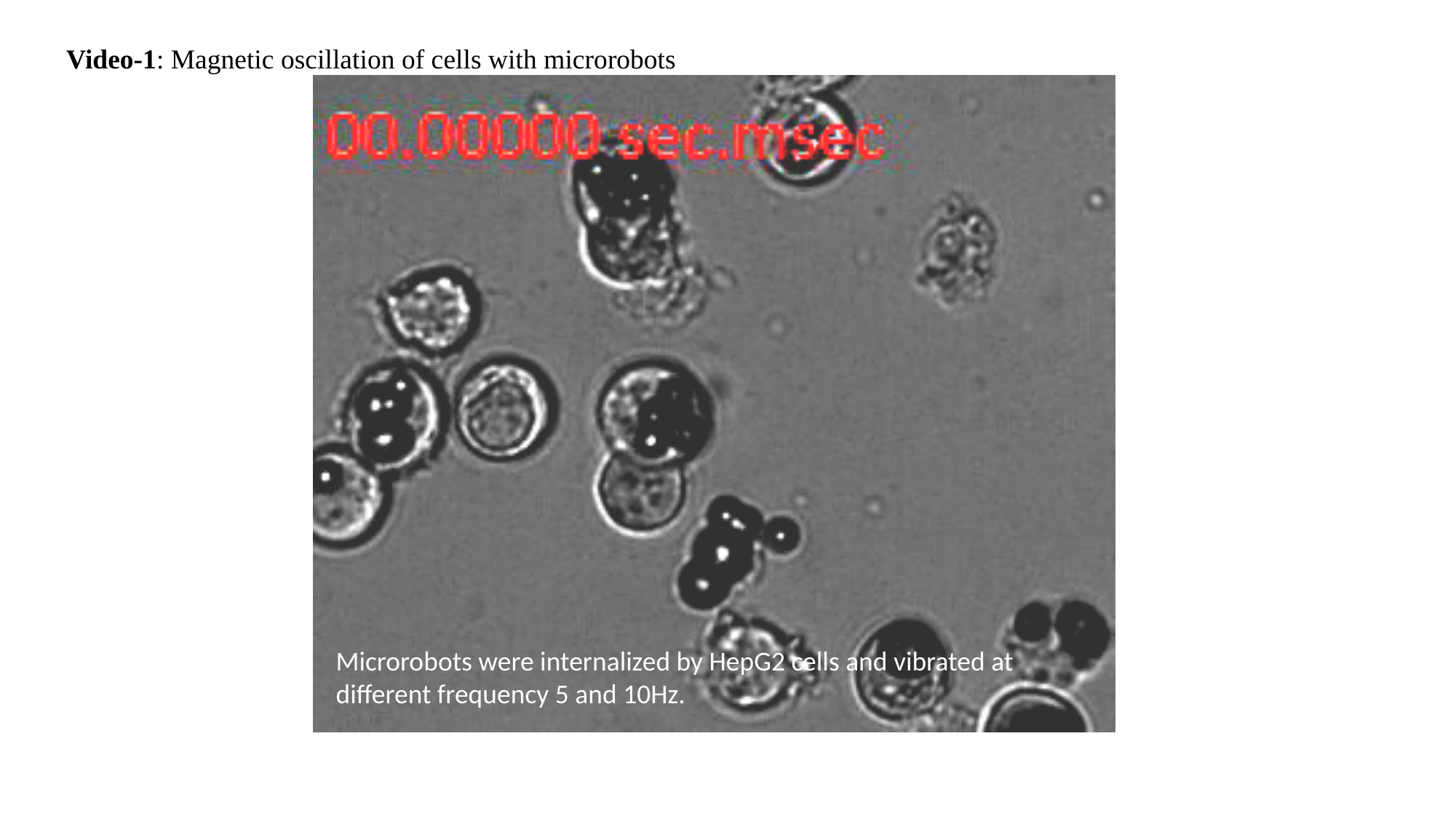

Video-1: Magnetic oscillation of cells with microrobots
Microrobots were internalized by HepG2 cells and vibrated at different frequency 5 and 10Hz.
